# Activating Cognitive Processes in Human Cortex at Multiple Frequency Bands

**DOI:** 10.1101/2024.07.26.605234

**Authors:** Zhu-Qing Gong, Xi-Nian Zuo

**Affiliations:** State Key Laboratory of Cognitive Neuroscience and Learning, Beijing Normal University, Beijing 100875, China; Developmental Population Neuroscience Center, IDG/McGovern Institute for Brain Research, Beijing Normal University, Beijing 100875, China; National Basic Science Data Center, Beijing 100190, China; Institute of Psychology, Chinese Academy of Sciences, Beijing 100101, China

**Author notes:** Correspondence to Xi-Nian Zuo (;).

**Keywords:** task-fMRI, frequency, neural oscillation, cognitive functions

## Abstract

Neural oscillations are fundamental for brain function, governing various cognitive processes. While electrophysiological studies have characterized the frequency properties of these oscillations, their spatial resolution is limited. In contrast, functional magnetic resonance imaging (fMRI) provides high spatial resolution but was initially restricted to a narrow frequency range. This study aims to bridge the gap by investigating the role of blood-oxygen-level-dependent (BOLD) oscillations across multiple frequency bands in cognitive processes using high temporal resolution task-fMRI data. Our findings reveal that different frequency bands are associated with distinct functional processes. Specifically, the slow-1 to slow-3 bands primarily contribute to local sensory information processing, while the slow-4 band is crucial for various fundamental cognitive functions, including somatic motor function and social cognitive function. The slow-5 band is involved in cognitive processes requiring higher memory load, integrated cognitive processing, and attention maintenance. Through multiband activation analysis, this study underscores the importance of analyzing a broad frequency range to capture the full spectrum of brain function. These findings highlight the diverse roles of different frequency bands in brain activity, shedding light on the underlying mechanisms of cognitive processes. This research enhances our understanding of the neural mechanisms underlying cognitive processes and has significant implications for cognitive neuroscience and clinical applications.

## Introduction

Brain function is underpinned by various neural oscillations occurring at distinct frequencies. These oscillations govern different brain functions. Owing to their high temporal resolution, electrophysiological methods can record a wide range of high-frequency oscillations. Over the past century, electrophysiological studies have characterized the frequency properties of neural oscillations. In general, oscillations in higher frequency bands predominate in local and bottom-up functions, whereas oscillations in lower frequency bands are more prevalent in long-range communications and top-down functions (Buzsaki & Draguhn, 2004; Kopell et al., 2000). For example, sensory stimuli initially evoke higher-frequency gamma oscillations, followed by the observation of lower-frequency beta oscillations (Kopell et al., 2000). Similarly, in visual processing, local activities engage gamma oscillations, cross-cortex semantic processing engages beta oscillations, and long-range connections involve oscillations in a lower frequency range (theta or alpha) (von Stein & Sarnthein, 2000). These phenomena reflect the temporal structure involved in the processing of external stimuli. Higher-frequency oscillations mediate stimuli detection and local primary processing, and information transitions to lower-frequency oscillations for further long-range integrative processing. The temporal structure of neural oscillations has significant implications for biological processes. A fast response is crucial for detecting external stimuli, especially dangerous signals. High-frequency oscillations that rapidly transmit information could satisfy this demand. In contrast, higher-order cognitive processing, such as learning, requires a longer processing window, facilitated by low-frequency oscillations. Additionally, the white matter structure provides the physiological basis for this frequency structure (Aboitiz, 1992). Axons in the sensory and motor cortex are thick and well-myelinated, allowing them to transfer neural activity very quickly. Associated regions and distant brain areas are connected by thin and sparsely myelinated axons that transfer information more slowly. This physiological basis ensures that the frequency characteristics of neural oscillations remain consistent regardless of measuring techniques. Therefore, neural oscillations recorded by other methods, such as functional magnetic resonance imaging (fMRI), are expected to exhibit frequency-specific properties and functions.

Although electrophysiological methods have high temporal resolution, their spatial resolution is not sufficient to map the distribution of neural oscillations across the brain. fMRI, which measures neural activity through high spatial resolution blood-oxygen-level-dependent (BOLD) signals, could fill this gap. Owing to the limited sampling rate and signal-to-noise ratio of early MRI devices, early resting-state fMRI studies focused only on a single frequency range (approximately 0.01 to 0.1 Hz), and most subsequent fMRI studies also used this conventional frequency range, thus ignoring the frequency information of the BOLD signal (Biswal et al., 1995). With the improved performance of MRI techniques, BOLD oscillations recorded with fast-sampling sequences can cover up to six frequency bands (e.g., slow-1, slow-2, … slow-6), making fMRI an excellent instrument for measuring the precise cortical distribution of neural oscillations across different frequencies. In the last decade, an increasing number of researchers have begun to investigate the frequency characteristics of BOLD oscillations. Both regional and network-level metrics commonly used in traditional single-frequency fMRI studies show frequency dependency (Jamadar et al., 2018; Li et al., 2018; Ma et al., 2021; Ries et al., 2018; Xue et al., 2014). Low-frequency BOLD oscillations are more active in high-order association areas, and high-frequency BOLD oscillations are more active in primary sensory and motor regions (Baria et al., 2011; Gong et al., 2021). At low frequencies, BOLD oscillations demonstrate more and stronger long-range connections, and networks exhibit greater global efficiency (Park et al., 2019; Thompson & Fransson, 2015; Yan et al., 2009). However, at higher bands, connections are more localized and exhibit greater network efficiency (Park et al., 2019; Thompson & Fransson, 2015). These findings reveal that BOLD oscillations have similar frequency characteristics to those of electrophysiological oscillations, attributable to the same physiological entity of neural oscillations being recorded by different techniques.

There is still a missing link between the frequency characteristics of BOLD oscillations and the cognitive functions involved with each frequency band of BOLD oscillations. How BOLD oscillations are organized in the frequency domain and how BOLD oscillations in different frequency bands are activated during cognitive processes remain major questions. In attempting to fill this gap, we recently investigated the frequency hierarchy of the six frequency bands of intrinsic BOLD oscillations (Gong & Zuo, 2023). We found that different brain networks demonstrate the highest integration level in different frequency bands, indicating that brain networks integrate information at different speeds. Generally, primary sensory and motor regions integrate information more rapidly than association regions. Additionally, meta-analytic decoding revealed that from slow-1 to slow-6, the cognitive functions associated with brain regions showing the highest level of integration transitioned from primary sensory and perception functions to more complex cognitive processing. These findings preliminarily reveal the brain functions embedded in the intrinsic BOLD oscillations. The next step is to provide concrete evidence to demonstrate the functions involved in each frequency band. Task-fMRI is a direct method for locating brain regions activated by cognitive tasks. By analyzing task activation across multiple frequency bands, it is also possible to locate the frequency bands activated by tasks. However, similar to most resting-state fMRI studies, previous task-fMRI studies were conducted in a single frequency range; thus, we cannot infer which frequency band brain regions are activated. In the present study, we plan to conduct multiband task activation analysis on seven tasks to provide substantial evidence of the functions involved in different frequency bands. According to findings from multiband resting-state studies, we aim to test the following hypotheses. First, cognitive processes are coordinated by oscillations of multiple frequency bands; thus, each task will activate brain regions in more than one band. Second, each band governs distinct functions; thus, for a single task, the spatial activation patterns differ across frequency bands. The stimuli in each task may activate sensory regions in higher frequency bands, while further task processing may activate regions in lower frequency bands. Third, task processing that requires more cognitive load will activate lower bands than tasks that require less cognitive load.

## Methods

### Data acquisition and preprocessing

We used task-fMRI data from the Human Connectome Project (HCP) S1200 young adult dataset in this study because the HCP data were acquired with high temporal resolution and the task-fMRI data included seven tasks; thus, we could explore the distribution of multiple functions in multiple frequency bands. We included data from 339 unrelated participants in our analysis. All participants completed seven task-state scans, including working memory (N-back task), incentive processing (gambling task), motor mapping, language processing (story task), social cognition (theory of mind task), relational processing (dimensional change detection task), and emotion processing (Hariri task). All task scans were block-designed, and each task scan comprised two runs. Details of the task paradigms and scan parameters are described in (Barch et al., 2013). The data were preprocessed according to the HCP minimal preprocessing pipeline with the ICA-FIX denoising procedure (Glasser et al., 2013). The preprocessing procedures included (i) gradient distortion correction; (ii) motion correction; (iii) EPI image distortion correction; (iv) registration to the T1w image; (v) one-step spline resampling; (vi) intensity normalization and brain masking; (vii) transfer of volume-based time series into surface-based time series; and (vii) ICA-FIX denoising.

### Frequency decomposing

The frequency band distribution of neural oscillations follows the natural logarithm linear law (N3L) (Penttonen & Buzsáki, 2003). On the natural logarithm axis, the center of each frequency band falls on an integral point, and the bandwidth of each band equals 1. According to the N3L, BOLD oscillations can be segmented into multiple frequency bands. The specific number of segmented frequency bands and band boundaries is determined by the scanning parameters (scanning duration and sampling rate). We used the DREAM toolbox to segment the preprocessed data (Gong et al., 2021).

### Task activation analysis

All tasks contained at least two task conditions, usually an experimental condition (e.g., the face matching condition in the emotion processing task) and a control condition (e.g., the shape matching condition in the emotion processing task). To fully characterize the activation patterns of each frequency band under different experimental conditions, we not only analyzed the contrasts between the experimental condition and the control condition, but also between all conditions and the baseline. At the participant level, for each run of each task, an activation map was calculated for every frequency band using FSL (Jenkinson et al., 2012). Then, for the two runs of each task, fixed-effects analyses were performed using FEAT to estimate the average effects across runs for each frequency band. Finally, at the group level, for each frequency band of each task, mixed-effects analyses were conducted using FLAME to estimate the average effects across participants. The group maps were visualized using Connectome Workbench with a threshold of z = 4.873 (Bonferroni-corrected p < 0.05).

## Results

### Frequency decomposition results

According to the scanning parameters, DREAM decomposed the BOLD oscillations of each task into bands ranging from slow-1 to slow-5. The sampling rate of all tasks was 0.72 seconds, so the highest frequency bands covered part of the slow-1 band. Since the scanning durations of all tasks differed, the lowest detectable frequencies varied among tasks. Notably, according to Nyquist’s theorem, the lowest frequency band of three tasks (the working memory task, language processing task, and motor task) could reach the slow-5 band. However, to guarantee the stationarity of the lowest frequency band, the DREAM toolbox discarded the four lowest frequency bins. After this adjustment, only the working memory task covered the slow-5 band. To reveal more comprehensive multiband activation characteristics, we still analyzed the slow-5 data for the language and motor tasks. The detailed boundaries of the decomposed frequency bands for the seven tasks are listed in Table 1.

**Table 1.** boundaries of the decomposed frequency bands for the seven tasks.

|  | slow-1 (Hz) | slow-2 (Hz) | slow-3 (Hz) | slow-4 (Hz) | slow-5 (Hz) |
| --- | --- | --- | --- | --- | --- |
| working memory | 0.6055-0.6944 | 0.2224-0.6055 | 0.0821-0.2224 | 0.0308-0.0821 | 0.0205-0.0308 |
| language processing | 0.6065-0.6944 | 0.2242-0.6065 | 0.0835-0.2242 | 0.0308-0.0835 | 0.0264-0.0308 |
| relational processing | 0.6046-0.6944 | 0.2215-0.6046 | 0.0838-0.2215 | 0.0359-0.0838 |  |
| emotion processing | 0.6076-0.6944 | 0.2210-0.6076 | 0.0789-0.2210 | 0.0473-0.0789 |  |
| gambling | 0.6070-0.6944 | 0.2242-0.6070 | 0.0820-0.2242 | 0.0328-0.0820 |  |
| motor | 0.6064-0.6944 | 0.2250-0.6064 | 0.0831-0.2250 | 0.0293-0.0831 | 0.0205-0.0293 |
| social cognition | 0.6083-0.6944 | 0.2230-0.6083 | 0.0811-0.2230 | 0.0304-0.0811 |  |

### Working memory

The working memory task data could be decomposed into five frequency bands: from slow-1 to slow-5. The working memory task contained two conditions: 0-back and 2-back. Both conditions’ activation patterns showed spatial separation across frequency bands (Figure 1). In the slow-1 to slow-3 bands, the 0-back and 2-back conditions both activated the visual cortex. In slow-1, the activation region was constrained in the primary visual cortex. In slow-2, the activation area extended and covered the secondary visual cortex. In slow-3, the regions activated were concentrated in the ventral visual cortex. The activation of the visual cortex in these high-frequency bands is consistent with our hypothesis that the detection and local processing of sensory stimuli occurs in the high-frequency bands. In the slow-4 band, both conditions activated large areas in the visual cortex and the association cortices including the middle frontal gyrus (control network), the anterior insula (ventral attention network), the dorsal anterior cingulate cortex (ventral attention network), the superior parietal lobule (dorsal attention network), and the middle temporal gyrus (language network). The 0-back condition exhibited greater spatial extension than the 2-back condition in this band, especially in the occipital and temporal lobes. In the slow-5 band, the activation patterns of the two conditions were similar to those in the slow-4 band but with greater spatial extension in the frontal and parietal lobes and smaller activation areas in the occipital and temporal lobes. Although the activation patterns of the 0-back and 2-back conditions were similar across frequency bands, the contrast between conditions revealed that the higher memory-load function might be processed in the slow-5 band. The 2-back vs. 0-back contrast showed significant activation only in the slow-5 band. The activated areas were mainly localized in the control network, the ventral attention network, and the medial occipital cortex. Meanwhile, the 0-back vs. 2-back condition significantly activated the slow-3 to slow-5 bands. In particular, in the slow-4 band, the activation was the most robust and widespread across the cortex. The contrast in activation between the two conditions implied that slow-4 was more involved in basic cognitive processing with low memory load, while slow-5 was more involved with complex cognitive processing with high memory load.

**Figure 1.**
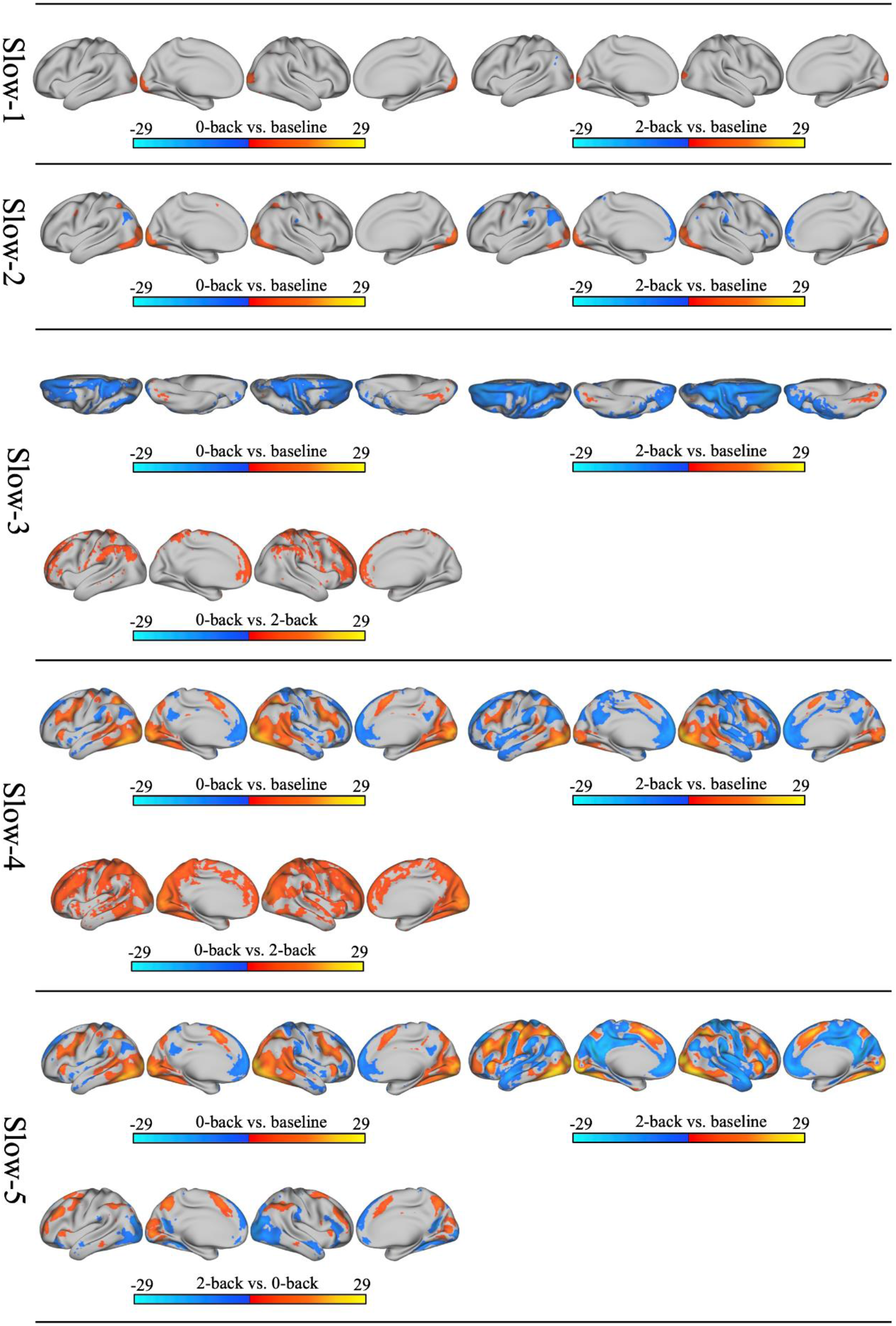
The multiband activation maps of the working memory task.

### Language processing

The language processing task contained two conditions: math and story. Auditory stimuli were used in both task conditions. The BOLD oscillations of the language processing task were decomposed into slow-1 to slow-5 frequency bands. The multiband activation patterns are shown in Figure 2. In the slow-1 and slow-2 bands, both conditions elicited significant activation only in the superior temporal cortex. In slow-4, in addition to robust activation in the auditory cortex, both conditions also activated regions in the lateral frontal cortex (the control network), the dorsal anterior cingulate cortex (the ventral attention network), and regions in the visual network. In slow-5, the activation areas expanded for both conditions, indicating more brain regions participated in information processing. It can be inferred that the local processing of auditory stimuli occurred in high-frequency bands (slow-1 and slow-2), while further integration occurred in the lower band (slow-4 and slow-5). The story vs. math contrast activated the bilateral inferior parietal lobule (the default mode network), the superior frontal gyrus (the control network), and the medial prefrontal cortex (the default mode network) in the slow-3 band, and regions in the default mode network, the language network, the somatomotor network, and the visual network in slow-4 and slow-5. Therefore, we infer that semantic processing mainly involves the slow-3 to slow-5 band. Particularly, the activation in multiple primary sensory cortices in slow-4 and slow-5 indicates that deep semantic comprehension emerges in lower frequency bands and triggers sensory representations (such as visual representations related to the content of the story). The math vs. story contrast only elicited significant activation in the slow-4 and slow-5 band. In slow-4, the activation regions were localized in the control network and the dorsal attention network, especially the lateral side of the cortex. In slow-5, activation regions in the control network expanded, and large regions in the ventral attention network and medial visual network were activated. Therefore, we infer that arithmetic function is mainly associated with the slow-4 and slow-5 bands.

**Figure 2.**
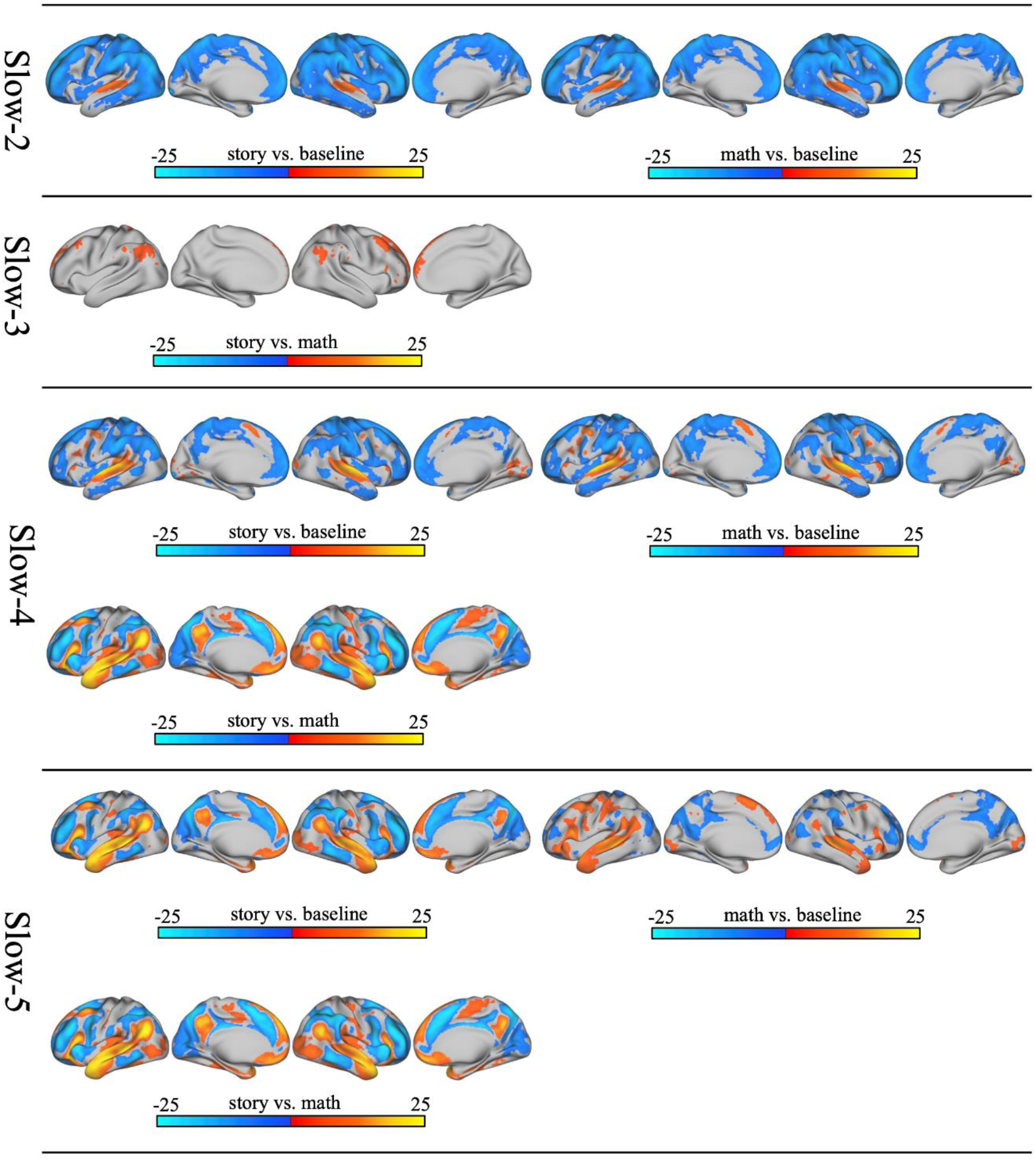
The multiband activation maps of the language processing task.

### Relational processing

The relational processing task data were decomposed into four frequency bands (slow-1 to slow-4). The task contained two conditions: a relational processing condition and a control match condition. According to previous single-frequency range studies, the relational vs. match condition can reliably activate the anterior prefrontal cortex. The multiple-band activation results showed that the relational vs. match condition significantly activated large areas in the anterior prefrontal cortex only in the slow-4 band (Figure 3). Thus, relational processing may occur in the slow-4 band. In addition, both conditions activated the visual cortex and the intraparietal sulcus in the slow-3 band. The match vs. relational condition activated sparse regions in the posterior parietal cortex (the control network) in slow-4.

**Figure 3.**
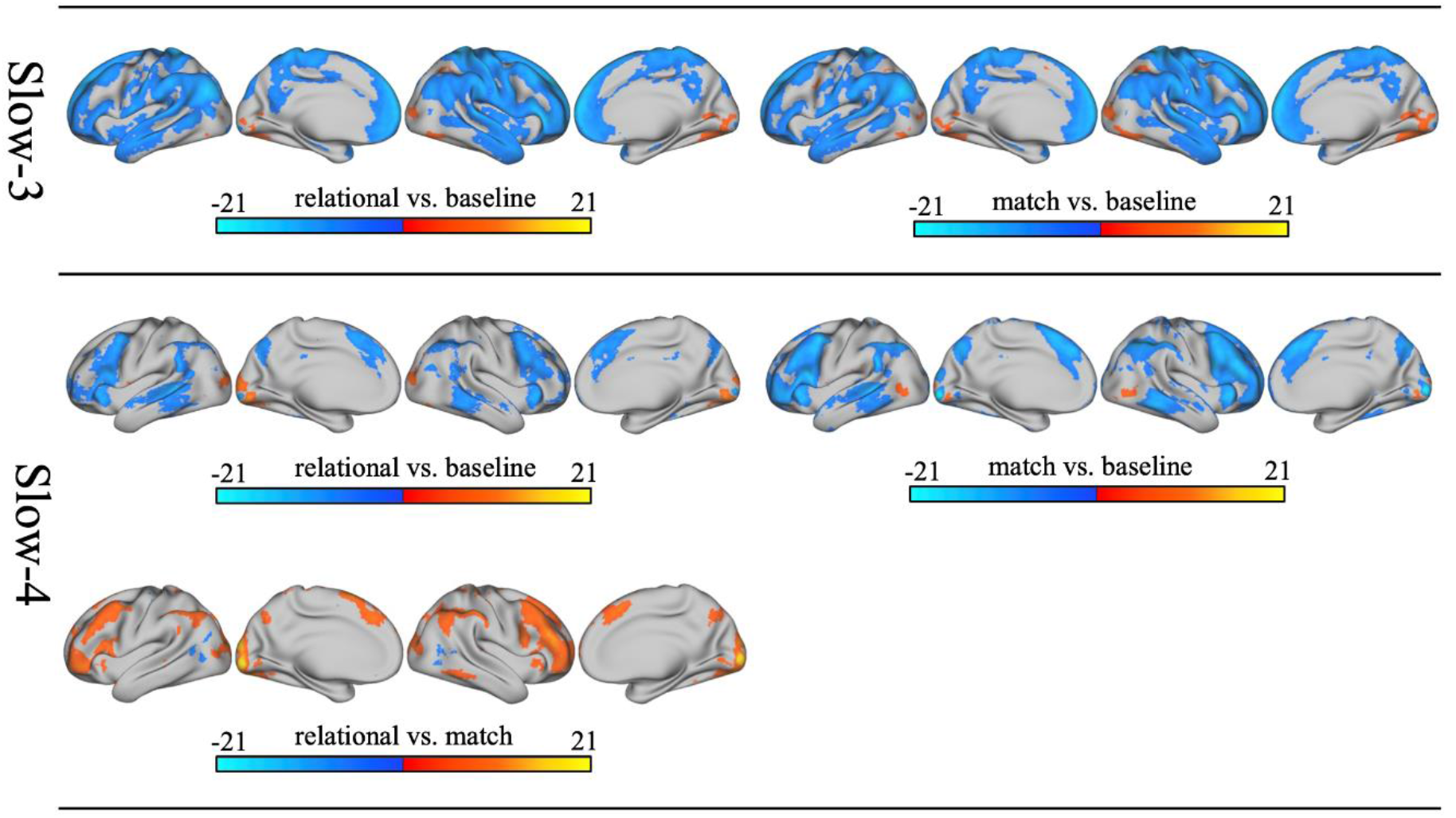
The multiband activation maps of the relational processing task.

### Emotion processing

This task contained a face-matching condition and a control condition of shape-matching. The emotion processing task data were decomposed into four frequency bands (slow-1 to slow-4). The multiband activation patterns are shown in Figure 4. The slow-1 band did not show any significant activation. In the slow-2 band, both the face condition and the face vs. shape condition activated small regions in the primary visual cortex. In slow-3, both conditions activated the occipital cortex (the visual network) and the superior parietal lobule (the dorsal attention network). The face vs. shape contrast activated the ventral lateral occipital cortex, especially the right side, in the slow-3 band. In the slow-4 band, the face vs. shape contrast elicited activation in the dorsolateral prefrontal cortex (the control network and dorsal attention network), the superior parietal lobule (the dorsal attention network), and the occipital cortex (the visual network). Therefore, we infer that the processing associated with emotional faces involves BOLD oscillations in the slow-4 band. Additionally, the shape vs. face contrast activated large regions in the frontal and parietal cortex in the slow-3 band.

**Figure 4.**
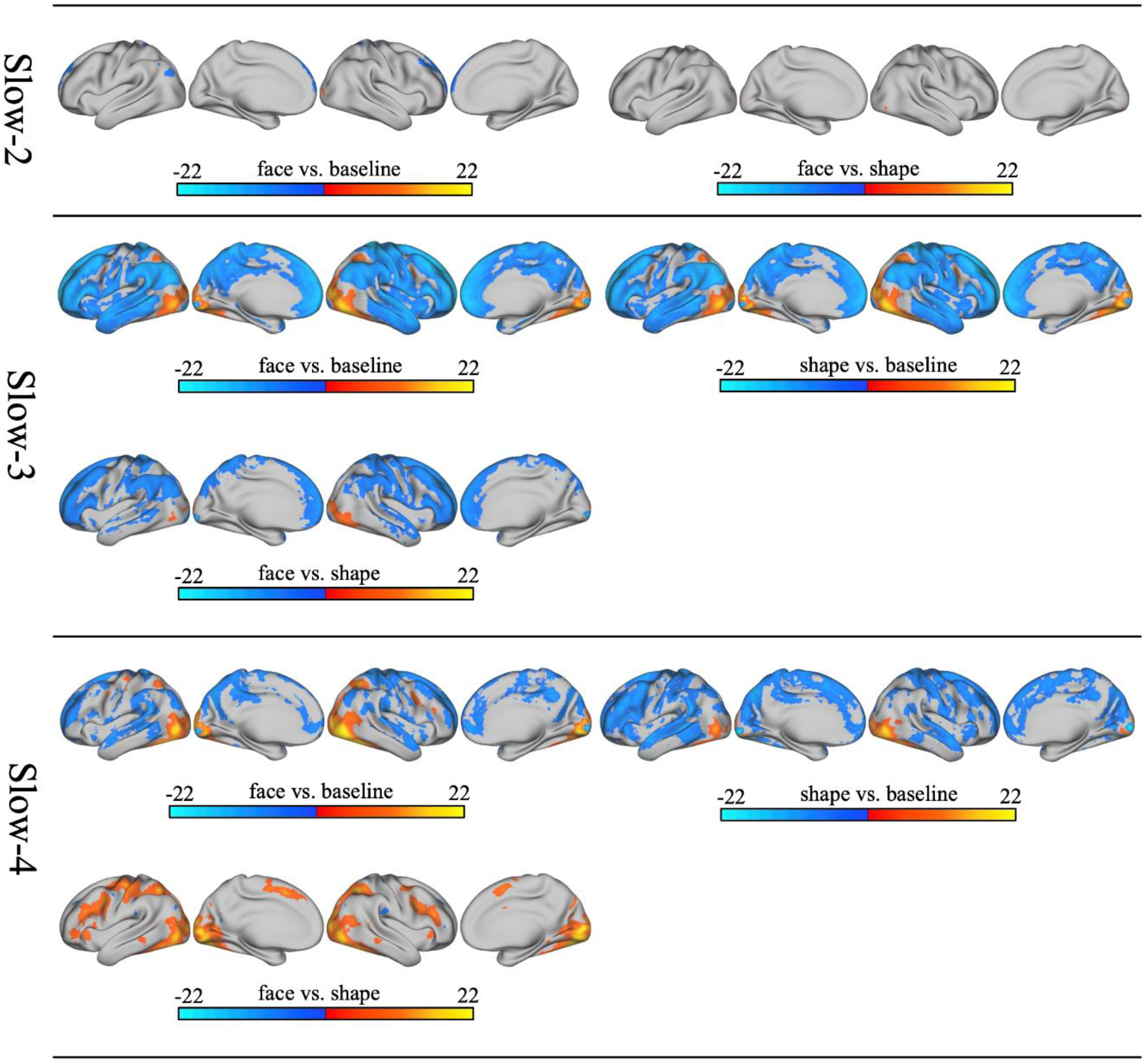
The multiband activation maps of the emotion processing task.

### Gambling

This task included a reward condition and a punishment condition. The frequency range of the data spans from slow-1 to slow-4. The multiband activation patterns are shown in Figure 5. Neither the slow-1 band nor the slow-2 band showed significant activation. In the slow-3 band, the punishment and reward conditions both activated small regions in the primary visual cortex. In addition, the reward condition also activated the left angular gyrus and small regions in the right somatosensory cortex. The reward vs. punishment contrast activated small regions in the bilateral primary visual cortex and the right somatosensory cortex. In the slow-4 band, both the reward and punishment conditions activated the ventrolateral occipital lobe, especially the right hemisphere. In addition, the punishment vs. reward condition activated the bilateral anterior insula (the control network and the ventral attention network), middle frontal cortex (the control network), superior parietal lobule (the dorsal attention network), dorsal anterior cingulate cortex (the control network), dorsolateral and ventromedial occipital cortex (the visual network), the left premotor cortex (dorsal attention network), and the left sensorimotor cortex (somatomotor network). The reward vs. punishment contrast only activated a small region in the primary visual cortex in slow-4. One possible explanation for these results is that negative emotions induced by the punishment condition may increase the consumption of low-frequency cognitive and attentional resources while performing the same cognitive activities.

**Figure 5.**
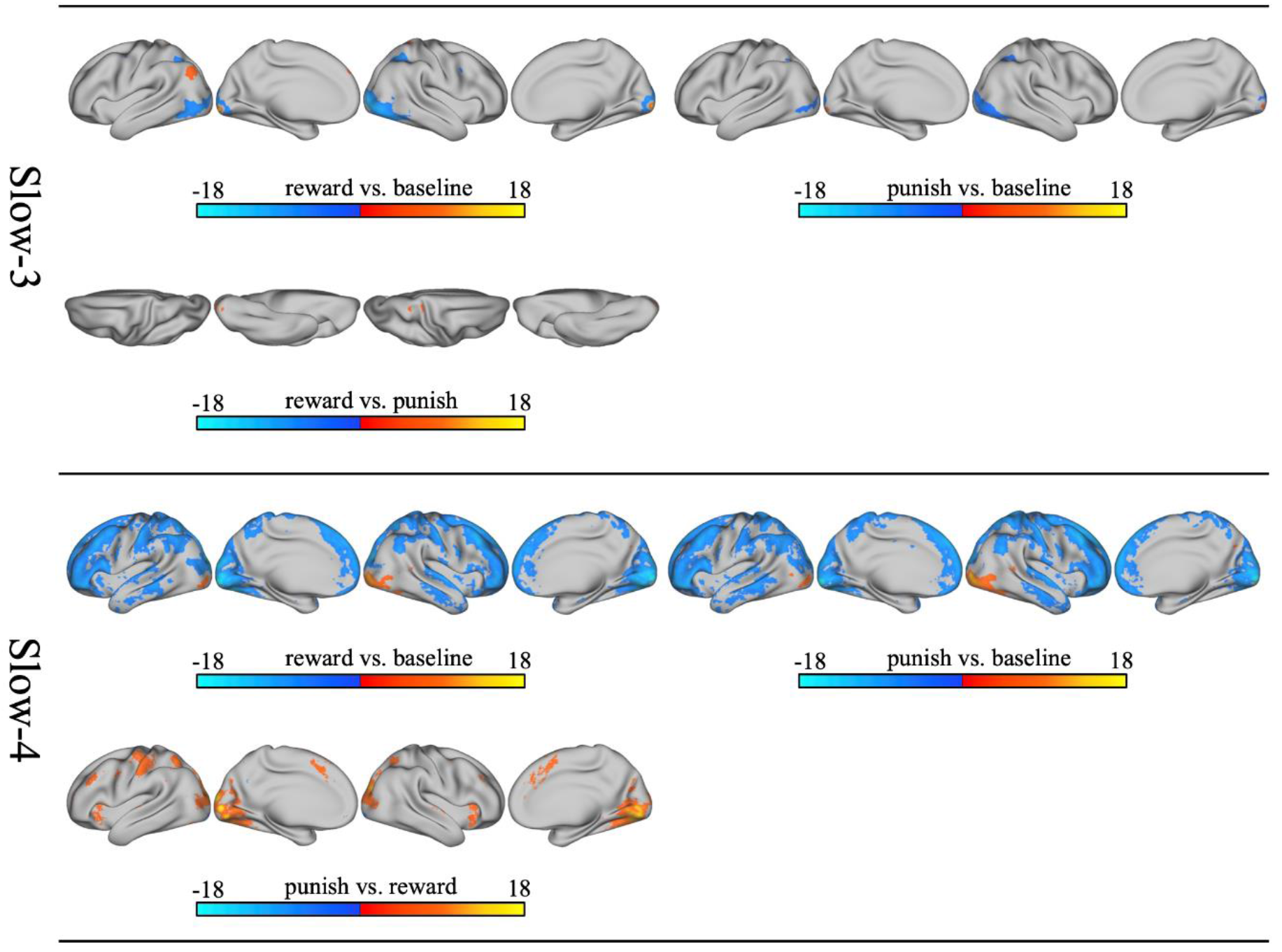
The multiband activation maps of the gambling task.

### Motor

This task contained five body-part movement conditions: tongue, left hand, right hand, left foot, and right foot. In addition to the motor conditions, we also analyzed the visual cue period, which was a 3-second duration of cue words prior to each movement block. The motor task timeseries was decomposed into five frequency bands (slow-1 to slow-5). The multiband activation patterns are shown in Figure 6. No significant activations were found for the slow-1 or slow-2 bands. The visual cue elicited significant activations in the occipital cortex (visual network), the dorsal cingulate cortex (ventral attention network), and the left premotor cortex (dorsal attention network) in the slow-3 band and in extensive brain regions other than the default mode network, the inferior and middle frontal gyrus, and parts of the motor cortex in the slow-4 band. In slow-3, the average of motor conditions activated the right angular gyrus (default mode network), bilateral medial prefrontal cortex (default mode network), and posterior medial occipital cortex (visual network). In slow-4, the motor conditions separately activated the motor cortex corresponding to the body parts in motion. The tongue condition activated the bilateral tongue regions in the sensorimotor cortex. The left- and right-foot conditions activated the contralateral foot regions in the sensorimotor cortex. The left- and right-hand conditions activated the contralateral hand regions in the sensorimotor cortex. In slow-5, the visual cue activated large areas outside the sensorimotor cortex, including the dorsal and ventral attention networks, the anterior insula (ventral attention network), the posterior temporal lobe (ventral and dorsal attention networks), the dorsal lateral prefrontal lobe (control network), and the medial visual network. In addition to activating the same regions as the visual cue, the motor conditions also activated the sensorimotor cortex corresponding to the body parts in motion.

**Figure 6.**
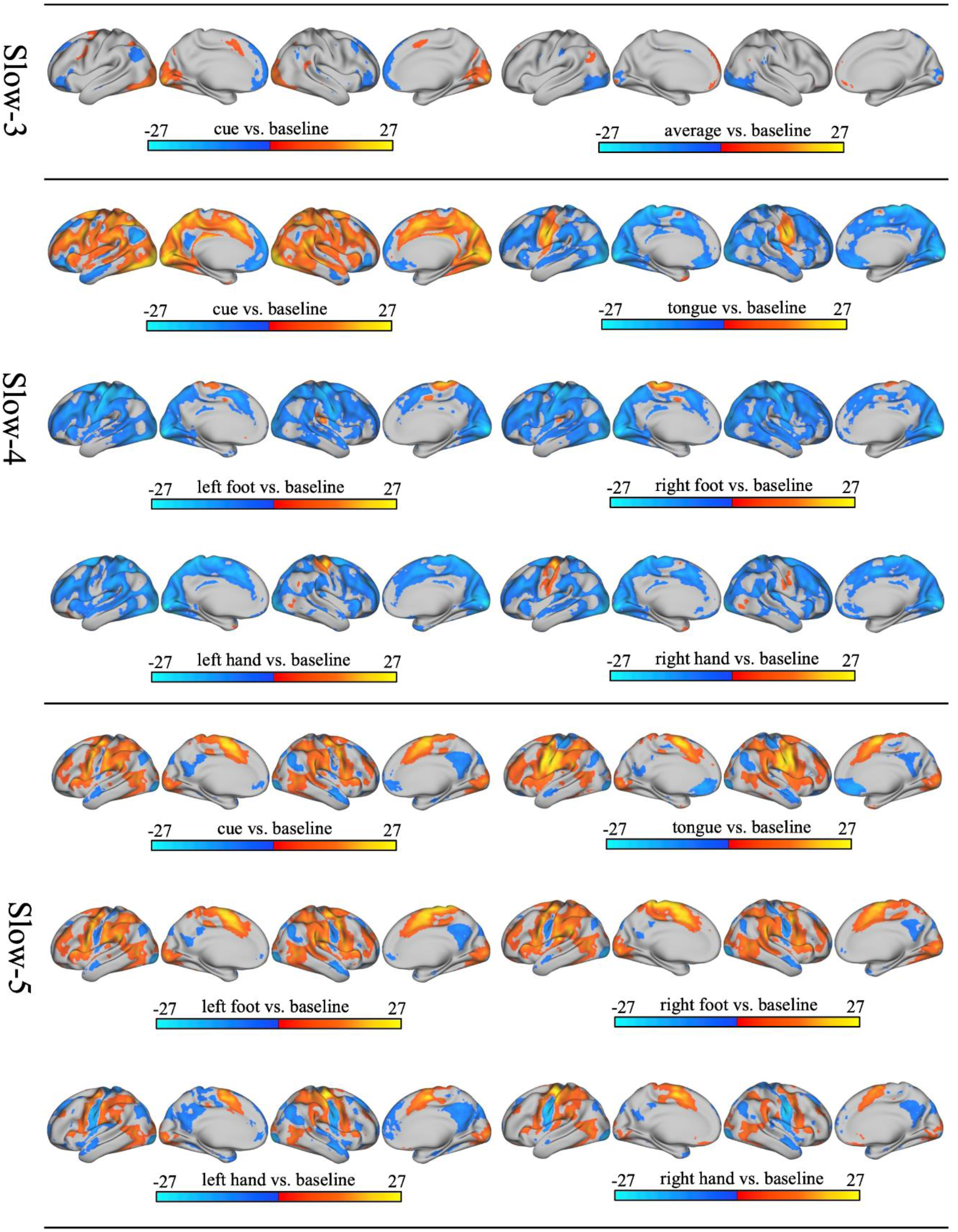
The multiband activation maps of the motor task.

### Social cognition

The social cognition task included a theory of mind (ToM) condition and a random control condition. The data from this task were decomposed into four frequency bands: slow-1 to slow-4. The activation patterns across these frequency bands are depicted in Figure 7.

**Figure 7.**
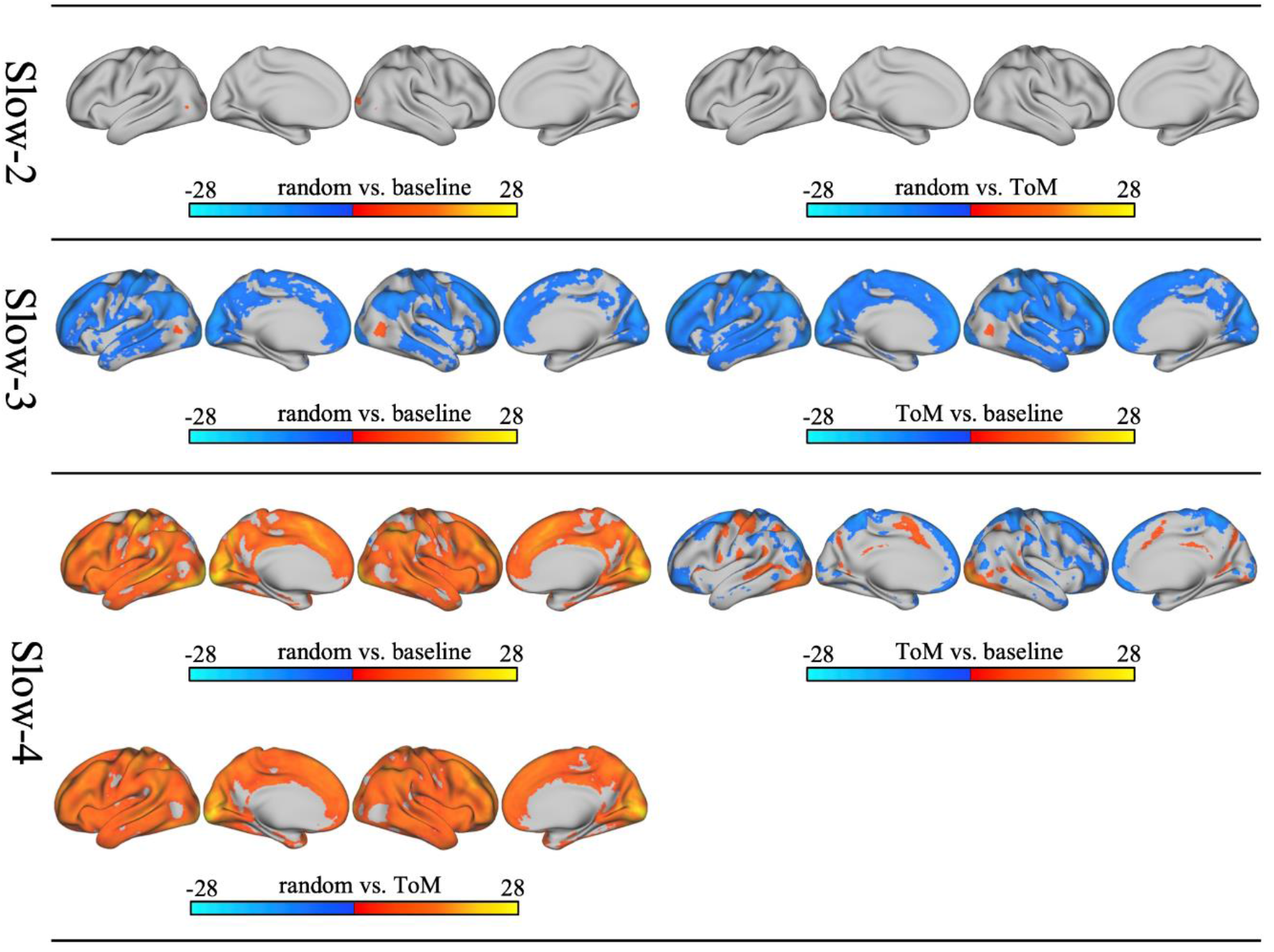
The multiband activation maps of the social cognition task.

No significant activations were observed in the slow-1 band for either condition. In the slow-2 band, the random and random vs. ToM conditions activated small regions in the visual cortex. In the slow-3 band, both the ToM and random conditions resulted in activations of the lateral occipital cortex. In the slow-4 band, the ToM condition exhibited significant activations in the lateral occipital cortex (visual network), superior temporal sulcus (language network), superior parietal lobule (dorsal attention network), dorsal cingulate cortex (ventral attention network and control network), anterior insula (ventral attention network), as well as regions in the somatomotor network. Additionally, the random condition and the comparison between the random and ToM conditions activated widespread regions, with the most robust activations observed in the visual network.

## Discussion

In the present study, we examined the activation patterns of several cognitive tasks across distinct frequency bands. As expected, each task elicited activations in multiple frequency bands, indicating the involvement of multiple BOLD oscillation frequencies during task processing. The spatial activation patterns for each task exhibited variations among frequency bands, particularly between higher frequencies (slow-1 to slow-3) and lower frequencies (slow-4 and slow-5). This observation suggests that different frequency bands contribute uniquely to information processing and provides preliminary evidence that BOLD oscillations in distinct frequency bands govern specific functions.

### Slow-1 to slow-3: Detection and preliminary processing of visual and auditory information

While six of the seven task paradigms analyzed in this study (working memory, gambling, motor, social cognitive, relational processing, and emotion processing tasks) utilized visual stimuli to present experimental stimuli or task cues, one task (language processing task) employed auditory stimuli. Analysis of multiband task activation revealed that brain activity triggered by the physical properties of stimuli predominantly occurred in the slow-1 to slow-3 frequency bands. In these high-frequency bands, activation elicited by tasks presented with visual stimuli was mostly confined to the visual cortex, while activation induced by the auditory task was limited to the auditory cortex. Thus, activation of the visual or auditory cortex at higher frequency bands was determined by the physical properties of the stimuli (visual or auditory). Compared to association regions, the sensory cortex is characterized by thicker and more myelinated white matter fibers, enabling faster transmission rates (i.e., higher frequency of activity) (Aboitiz, 1992). This aligns with the observation that in high-frequency bands, task stimuli elicited activations mainly in the sensory cortex. Furthermore, in the slow-1 and slow-2 bands, visual and auditory stimuli exclusively activated their respective sensory cortices, suggesting that the processing of visual or auditory information is primarily confined to these areas within these two bands. In the slow-3 band, in addition to activation in the sensory cortex, certain tasks also elicited activation in small regions within the dorsal or ventral attention networks, indicating further integration of sensory information within this frequency range. For instance, in the relational processing task, both conditions activated the visual cortex and the intraparietal sulcus. The intraparietal sulcus is known to be involved in functions related to visual perception and attention, and it plays a crucial role in processing symbolic or numerical information, as well as visuospatial working memory (Artemenko et al., 2020; Grefkes & Fink, 2005). The simultaneous activation of the intraparietal sulcus and visual cortex in the slow-3 band suggests that the features of the visual stimuli have been further integrated and processed. In the motor task, the visual cue triggered activation not only in the visual cortex but also in the anterior dorsal cingulate cortex and the left premotor cortex within the slow-3 band. The white matter fibers in the premotor cortex directly project to the spinal cord, playing roles in motion-related functions and movement planning (Cisek & Kalaska, 2005; Rizzolatti & Sinigaglia, 2010). The concurrent activation of the premotor cortex and visual cortex in the slow-3 band suggests that the visual cue may have undergone semantic processing within this frequency range and synchronized with the premotor cortex for motor preparation. In summary, the processing of sensory stimuli in the slow-3 band extends beyond the sensory cortex and begins integrating functions across different brain regions. Therefore, we infer that the slow-1 to slow-3 bands primarily contribute to the preliminary processing of sensory information, with a hierarchical structure in sensory information processing: the slow-1 and slow-2 bands are responsible for the detection and local processing of sensory stimuli, while the slow-3 band is involved in the integrative processing of sensory information

This perspective is further supported by findings from a study on auditory perception (Frühholz et al., 2020). In this study, researchers identified several independent components in the auditory cortex during an auditory task. Within the slow-1 and slow-2 bands, a component emerged immediately after auditory stimuli and showed no preference for vocal or nonvocal stimuli in each band. In the slow-3 band, a component appearing later after the presentation of auditory stimuli exhibited a significant preference for vocal stimuli. The timing of component emergence may reflect the downward flow of information processing from higher frequencies, and the preference for vocal stimuli suggests a more integrated processing of auditory stimuli in the slow-3 band. Furthermore, the language processing task results showed that the story vs. math condition activated the angular gyrus in the slow-3 band, providing additional evidence that preliminary semantic processing may involve slow-3 oscillations.

The activity characteristics of BOLD oscillations at rest also support the role of higher frequencies in the detection and preliminary processing of sensory stimuli. Specifically, the salience network exhibits the highest power values in the high-frequency bands compared to other brain networks, while the auditory and visual cortices also show higher ALFF values in these bands (Gohel & Biswal, 2015; Gong et al., 2021). Moreover, the sensory cortices exhibit the highest levels of integration within the slow-1 to slow-3 bands (Gong & Zuo, 2023). These findings suggest that, even during the resting state, the brain regions responsible for salience detection and sensory processing are most active at higher frequencies.

### Slow-4: Motor function, basic cognition and modular segregation

During the resting state, most networks, excluding the default mode network and the visual network, exhibit higher power values in the slow-4 band compared to other frequency bands (Gohel & Biswal, 2015). The basal ganglia, a region related to motor functions, also demonstrates the highest ALFF values in the slow-4 band (Zuo et al., 2010). Additionally, networks in the slow-4 band show higher levels of segregation and network efficiency, and lower levels of global integration, suggesting that brain function is organized in a more modular manner within this band (Thompson & Fransson, 2015). Therefore, oscillations in the slow-4 band may be responsible for various fundamental functions, or functions with low demand for global integration, by facilitating modular segregation.

This perspective is supported by the activation patterns observed in multiple tasks within the slow-4 band. In the motor task, different motor conditions exclusively activated the corresponding motor cortex in the slow-4 band, indicating that the ability to control the movement of different body parts individually may be primarily governed by slow-4 oscillations. Furthermore, the working memory task exhibited more refined activation patterns in the slow-4 band, while activation in higher frequencies was limited to the visual cortex and was much broader in the slow-5 band. Both the 0-back and 2-back conditions activated the occipital cortex, the posterior middle frontal gyrus of the control network, the superior parietal lobule, the intraparietal sulcus of the dorsal attention network, and the middle temporal gyrus of the language network. The middle frontal cortex, superior parietal lobule, and intraparietal sulcus are brain regions associated with cognitive control (Ansari & Karmiloff-Smith, 2002; Babiloni et al., 2005; Fox et al., 2006; Koenigs et al., 2009). The concurrent activation of these brain regions suggests that working memory processes are predominantly mediated in the slow-4 band. Moreover, the 0-back vs. 2-back condition showed widespread activation in the slow-4 band. This suggests that neural activity in this frequency band is more involved in simple or basic cognitive processing.

In the language processing task, the contrast between math and story conditions showed significant activation in the slow-4 band. The math condition predominantly activated brain regions related to cognitive control, such as the dorsal lateral prefrontal cortex, posterior middle frontal gyrus, precentral gyrus, supramarginal gyrus, intraparietal sulcus, and regions in the temporal lobe. On the other hand, the story condition primarily activated brain regions associated with syntax and semantics, the sensory cortex, and the default mode network. The activation of the sensory cortex and default mode network may indicate that during story comprehension, participants generated sensory representations and recalled self-related memories or experiences. The contrast activation between the math and story conditions was more refined in slow-4 than in slow-5, indicating that the elementary processes of arithmetic and language comprehension are mainly mediated by slow-4 oscillations.

In the social cognition task, the social interaction video elicited activation in the superior temporal sulcus, a region related to the theory of mind, suggesting that social cognition is associated with slow-4 oscillations. In the relational processing task, the relational versus match condition activated regions in the lateral prefrontal cortex, inferior parietal lobule, right inferior temporal gyrus, and occipital cortex. Overall, the activation patterns in the relational processing, n-back, and math conditions are similar within the slow-4 band. These cognitive tasks all elicited activation in the lateral prefrontal cortex and lateral parietal cortex, consistent with previous findings highlighting the crucial roles of these regions in cognition. There are also variations in the precise location of these activation areas across tasks in this band, reflecting the modular segregation of functional organization during different cognitive tasks.

Based on the activation results of multiple tasks, it can be inferred that slow-4 is a key frequency band for intelligence. The BOLD oscillations within this frequency band play a crucial role in various cognitive functions, somatic motor function, and social cognitive function. This inference is also supported by a study that examined the frequency spectral properties of white matter signals, revealing that white matter voxels can be categorized into two groups based on their frequency spectrum density (Li et al., 2021). The first category consists of white matter voxels with a single peak in the frequency spectrum density, occurring at around 0.01Hz, close to the boundary between the slow-5 and slow-6 bands. The second category exhibits a double peak in the spectral density, with peak 1 occurring near 0.01Hz (similar to the single peak white matter voxels), and peak 2 occurring near 0.06Hz, close to the midpoint of the slow-4 band. The white matter complexity of voxels with dual peaks is higher, indicating the presence of different types of white matter fibers within these voxels: fibers with a signal transmission frequency of approximately 0.01Hz and fibers with a frequency of approximately 0.06Hz. The study further found that the power ratio of the two peaks (peak 2/peak 1) in dual-peak white matter voxels near the frontal lobe is positively correlated with cognitive task scores. The stronger the signal in the slow-4 band within these voxels, the higher the cognitive scores. This result provides confirmation of the role of the slow-4 band in cognitive function from the perspective of white matter fiber information transmission.

### Slow-5: function integration, memory load and attention

In the working memory task, the activation patterns of the two N-back conditions in the slow-5 band are similar to those in the slow-4 band. However, the activation range of the frontal and parietal lobes is more extensive in the slow-5 band, indicating that more brain regions are involved in task processing in this frequency band. Similarly, in the language processing task, the spatial activation range in the slow-5 band is wider than that in the slow-4 band. This suggests that the slow-5 band may play a role in more generalized and integrated cognitive processing compared to the modular segregation pattern observed in the slow-4 band. Intrinsic brain activity also reflects this characteristic, with different networks being densely connected in the slow-5 band while exhibiting segregation in the slow-4 band (Thompson & Fransson, 2015).

The slow-5 oscillations are also important for cognitive processes that require a high memory load. In the working memory task, the contrast of 2-back vs. 0-back only showed significant activation in the slow-5 band. The activated regions included the lateral prefrontal cortex, dorsal anterior cingulate cortex, precuneus, inferior parietal lobule, and medial occipital cortex. These activation regions, except for the precuneus, are similar to the activation regions observed in both conditions in the slow-4 band. This suggests that brain regions associated with working memory provide more computing resources for high-memory load tasks in the slow-5 band. The precuneus is associated with memory function and inhibitory responses in cognitive control (Boruchow & Hutchins, 1991; Lundstrom et al., 2005). It was only activated in the slow-5 band in the 2-back condition and 2-back vs. 0-back contrast, suggesting that its function in slow-5 is specifically related to memory processing. This finding is consistent with a recent study on resting-state brain networks, which identified the parietal memory network, with the precuneus as its core region, specifically in the slow-5 band (Luo et al., 2021). This further supports the notion that slow-5 oscillations are crucial for memory function. Overall, the activation patterns observed in the working memory task suggest that both the slow-4 and slow-5 bands play important roles in cognitive processing. The slow-4 band shows more modular segregation, while the slow-5 band exhibits more integrated and generalized activation patterns. We infer that slow-4 is more associated with elementary cognitive processing, while the slow-5 oscillations are more involved in high memory load processing. The specific activation of the precuneus in the slow-5 band further highlights the involvement of this frequency band in memory-related processes.

Attention maintenance may be another important function governed by slow-5 oscillations. Both the dorsal attention network and ventral attention network were largely activated in the slow-5 band. This phenomenon is particularly evident in the motor task. Unlike cognitive processing conditions, which exhibit similar spatial activation patterns in slow-4 and slow-5, all motor conditions showed distinct spatial activation patterns between the two bands. In slow-5, the motor conditions, as well as the visual cue, elicited large activations in the dorsal and ventral attention networks. This suggests that the attention-regulation function required to maintain task processes may be carried out through low-frequency oscillations. Additionally, these results align with resting-state meta-decoding findings that some regions showing the highest integration level in slow-5 are linked to attention function (Gong & Zuo, 2023).

### Limitations and future directions

In this study, we demonstrated that BOLD oscillations from multiple frequency bands participate in task processing and dominate distinct functions. This was revealed by the observation that various task conditions induce significant activations in multiple frequency bands, each with distinct spatial patterns. Therefore, to depict a comprehensive landscape of brain function, it is preferable to perform multiband frequency analysis in both resting-state and task-state studies. However, because the scanning durations of the task data we used were generally short (mostly less than 5 minutes), the slow-6 band could not be covered in this study, and not all tasks could cover the slow-5 band. Future studies should consider longer scan durations to cover more low-frequency ranges.

We computed the multiband activation maps using the conventional sustained response model, which consists of the hemodynamic response function convolved with a boxcar function following the experimental paradigm. The multiband activation maps of various tasks show clear differences between bands. In the slow-1 to slow-3 bands, the activated regions were primarily located in the visual or auditory cortex. In the slow-4 band, different task conditions evoked distinct sets of activated brain regions. The activated regions for each condition were spatially segregated but functionally associated, representing the organization of a set of modular network nodes. In the slow-5 band, the activated regions were more widespread and continuous, reflecting a more integrative organizational form. However, based on the response model we used, these activation maps reflect only brain areas with activity most consistent with the experimental paradigm design. Brain regions modulated by the task but oscillating in a different manner would not be modeled. For example, high-frequency oscillations in the association regions may be activated by task modulation, but the activation does not persist throughout the block and therefore does not show significant activation in the sustained response model analysis. Using model-free analysis, researchers have demonstrated that the whole brain is activated during a simple task (Gonzalez-Castillo et al., 2012). Future studies could use a more flexible response model to explore the functional mechanisms of multiband oscillations during tasks, especially in the higher frequencies.

Finally, this study provides a preliminary explanatory framework for multiband resting-state studies and clinical studies. For instance, a multiband study revealed that patients with Parkinson’s disease (PD) exhibited more altered brain activity in the slow-4 band (Hou et al., 2014). Furthermore, slow-4 demonstrated improved accuracy in distinguishing between PD patients and healthy controls as well as between PD patients with different symptoms (Hu et al., 2020; Tian et al., 2020). By confirming that the slow-4 is associated with motor function, it is reasonable to conclude that slow-4 is a key band for the dysfunction of PD patients. This framework not only advances our understanding of the multiband oscillatory mechanisms underlying cognitive and motor functions but also opens new avenues for clinical diagnostics and therapeutic strategies in neurological disorders. Future research should continue to explore these frequency bands to further elucidate their specific roles and potential applications in clinical settings.

## Notes

https://humanconnectome.org/

### Competing Interest Statement

The authors have declared no competing interest.

